# Agricultural production and CO2 emissions causes in the developing and developed countries: New insights from quantile regression and decomposition analysis

**DOI:** 10.1101/2020.11.16.384370

**Authors:** Rabnawaz Khan

## Abstract

Agriculture is the dominant economic activity of the economies. The developing and developed countries are responsible for the most greenhouse gasses emitted in the developing areas. Are there heterogeneous determinants of environmental degradation and CO2 emitters in developing and developed countries? and estimating the significance of agricultural production, renewable energy consumption, the industrial revolution, and economic growth. In this study, 22 countries’ environmental degradation analyze by two (per-capita and liquid) sources of CO2 emissions and using panel data from 1991 through 2016. This study adopts a panel regression (non-additive effects) and quantile regression techniques to explore the connection between agriculture and economic factors. And the extent of the CO2 emitter gap between developing and developed countries. The outcome of agriculture has a positive and significant influence on CO2 emission from liquid with a 36.75% increase in environmental degradation and a negative impact on CO2 emission in the total emissions by 19.12%. The agriculture-related activities negatively influence the environment, such as deforestation for feed cropping, burning of biomass, and deep soil cropping in the developing countries. Furthermore, the quantiles decomposition procedure in agriculture production is signifying heterogeneity of the determinants of environmental degradation, low and high CO2 emitters.

## 1. Introduction

The relationship between economic growth and CO2 emissions is, and will always remain, disputable. A few see the development of modern contamination issues, the need for success in managing worldwide warming, and the still rising populace within the Third World as confirmation positive that people are a short-sighted and avaricious species. However, like see the glass as half full. They note the more advance made in giving urban sanitation enhancements in the air quality and marvel at the continued improvement within the human condition made conceivable by innovative development in the developing and developed economies. The interface between economic growth and the environment has gotten much more consideration recently since the quickly extending empirical literature on the relationship between per capita income and contamination. This literature, known as the Environmental Kuznets Curve (EKC) approach that economic development initially leads to a deterioration in the environment hugely useful. So to a certain degree, the curve has presently turned, and distant less concern over the extreme fatigue of oil or magnesium, and distant more concern over air quality, worldwide warming, and the emanations of mechanical and industrial production [1]. Developing nations, with the fast advancement of the emerging economy, are driving the development of <u>energy</u> ingesting universally. The vitality was 7.645×109 (ton) of oil (toe) utilization was recorded in the developing nations, which was 58.2% all over the world in 2005. Additionally, the utilization of vitality was 2.37×109 (toe) increment in developing countries in 2015. The <u>energy</u> strength is 8.35 in China, 9.44 in Russia, and 3.87 in Germany, which shows an enormous hole between developing countries and developed countries. However, the emerging states diminish vitality gradually and attempt to attain bottleneck issues with well-developed innovation [2, 3].

Besides the industrial sectors, untenable agricultural practices due to bush burning, deforestation as well as biomass fuel burning can further determine the organic compound (oxidize soil) have been pointed out by 21% of the global share of GHG emissions. Agricultural practice and modification can naturally reduce CO2 emissions in the environment by soil organic compounds of the land cultivation [4]. Developing and developed the country’s investment in sustainable development will help to reduce building climate resilience and GHG emissions [5–7]. Fig.1 indicated that CO2 emissions approach of developing and developed countries by total CO2 emission and liquid fuel consumption.

**Figure 1:**
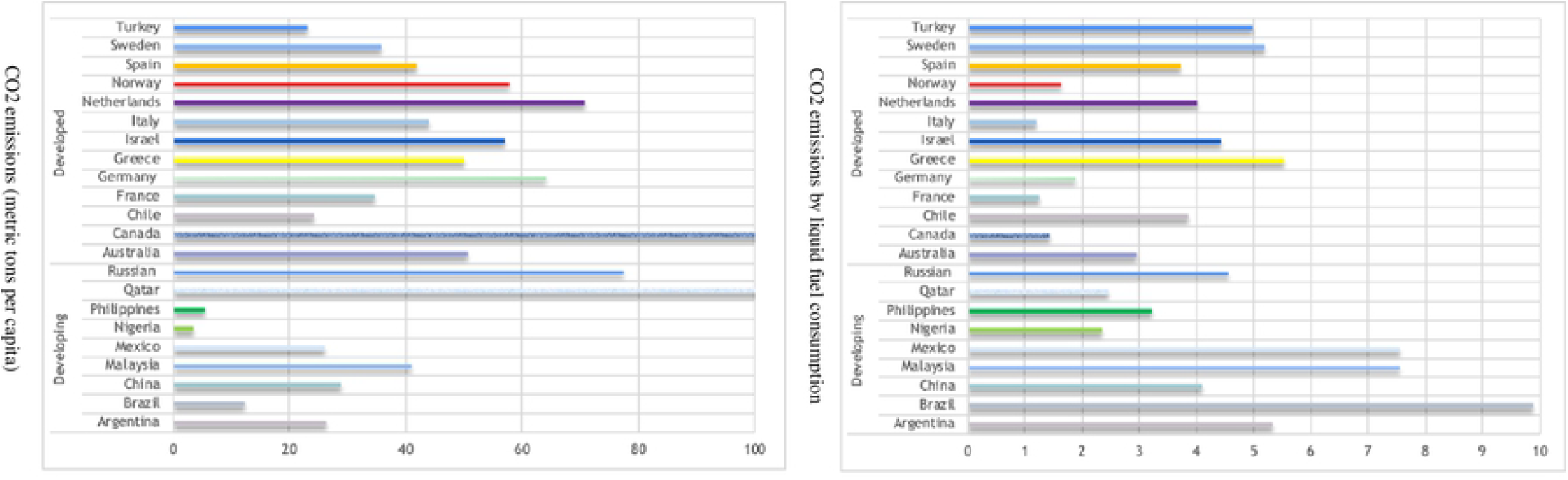
CO2 emissions of developing and developed countries. **Sources:** https://data.worldbank.org

CO2 emissions in developed (Nether land, Germany, and Canada) and the developing countries (Russia, Qatar, and Malaysia) are higher as well as the liquid consumption of developing (Brazil, Argentina, Mexico, and Malaysia) countries rapidly growing due to agriculture and industrial production. GHG emissions associated with agriculture totaled 582 million metric tons in 2017, which is slightly up from the prior year, but agriculture production down 2% from a decade ago. However, farmers contribute to CO2 emission confiscation efforts, i.e., forestry and wetlands, preservation of grasslands, greenhouse gas removal [8]. While improvement in agricultural productivity contributes to GHG climbed in 2017, it is also a substantial improvement in crop yield, nearly double the volume produced in developed countries in 1990. Overall, the implementation of technology in agriculture production, including animal genetics, crop, and chemical, has improved agriculture output without any significant inputs. The main objective of the present study is to determine the CO2 emissions consequence of agriculture production while directing for the income-induced EKC existing hypothesis as well as economic factors in the 22 countries. Our study of developing and developed economies is anticipated on the fact of developing countries as compared to developed countries agriculture productions and thus more economically integrated than developed countries.

<u>Agriculture</u> output and input comparisons for trends were made and use agricultural products in developing and developed countries. Agriculture production was measure across wheat, rice, maize, and other grains in different periods. The estimated results across all the commodities indicated that the growth rate of cereal production and yield significant slowdown previously Fig.2. Hence the analysis of the CO2 emission influence of agriculture productions rests on the inspection that agriculture is the most dominant factor of the economic sector in developing and developed countries. Fig.3 shows that the agriculture and cereal yield in developing and developed countries, and the results signified that China has high value-added agriculture, forestry, and fishing with cereal yield (kg per hectare) in the developing countries. Consequently, a production of 45% of agriculture production is a cause of 23% of C-LF. Also, Turkey has recorded the highest 6% production in AGR with 2% of C-LF and C-EM [9–11].

**Figure 2:**
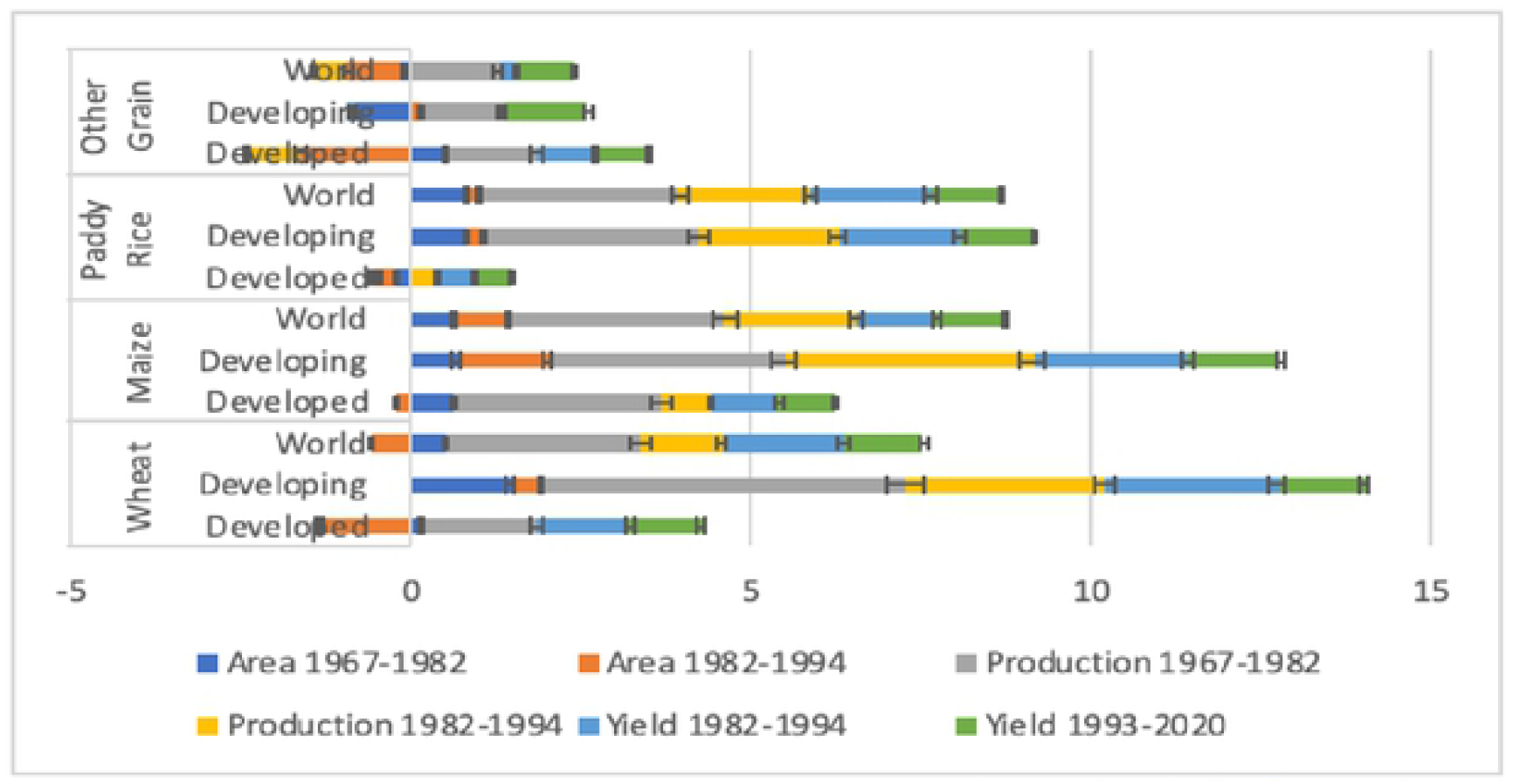
Agriculture productions. **Sources:** http://www.fao.org/3/x9447e07.htm#Notep:

**Figure 3:**
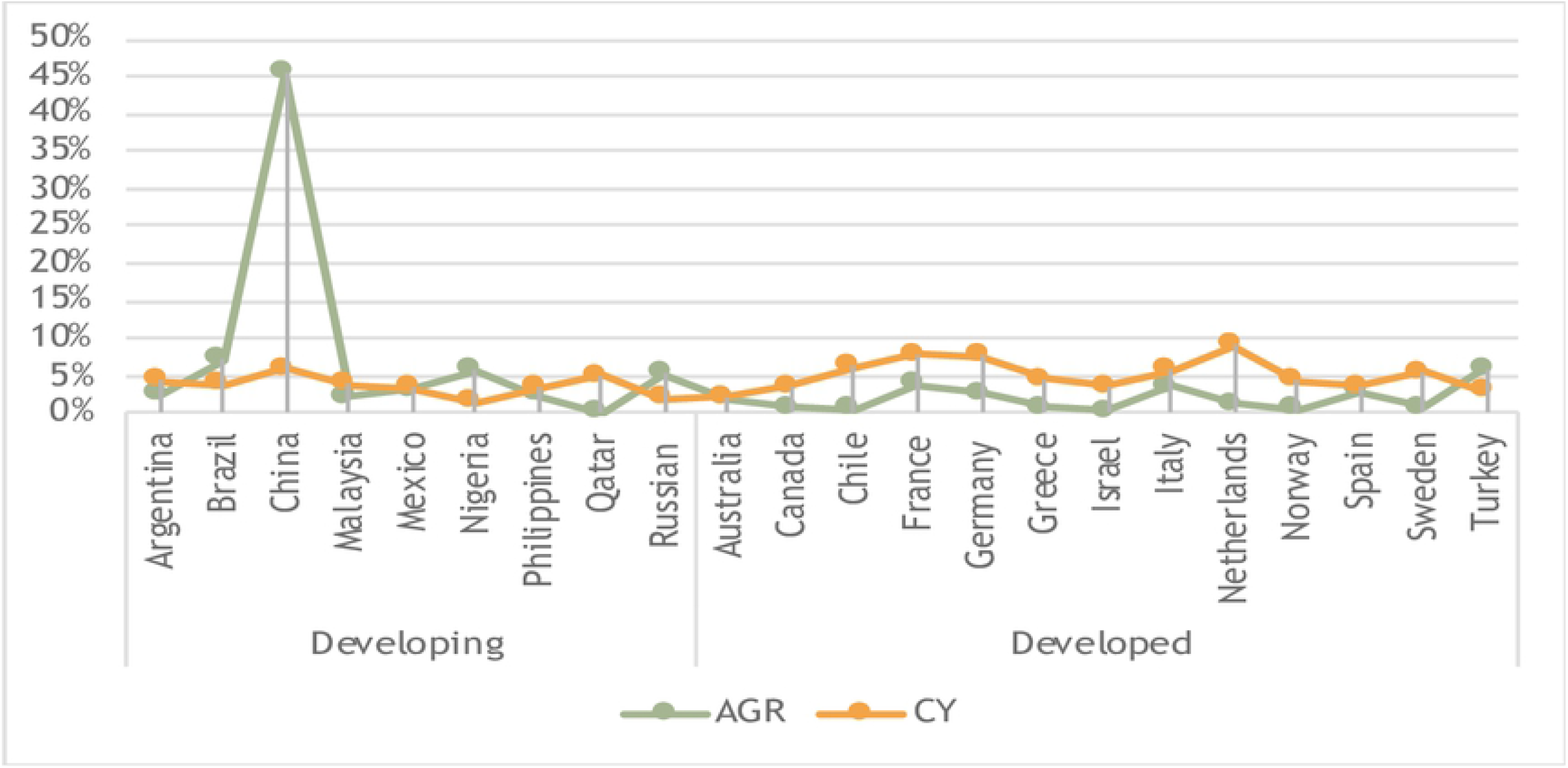
Carbon emissions with agriculture and cereal yield.

The EKC theory has been now investigated diverse thoughts in the carbon emanation. CO2 emissions and EKC reflect peak energy of intensity theory, says that energy increase in industrialization, then it reaches the peak, and finally decrease s. Hence the level of energy intensity shifted from the higher level and energy intensity of pollution-intensive industrial structure of the economy to the low intensity of light industries. The Sun’s argument implies that the carbon emissions EKC has only occurred in developing countries where an energy peak has occurred cause of abnormal economic and development growth [12, 13] [14]. Most parts of the study identified the explanatory research design, and the focus of the research is to provide the answers to why the agricultural production, economic growth, and development the U-shaped of EKC existing hypothesis. However, existing among the sources of income and ecological degradation, it inter-reconnect whenever increment increases and declines afterward level of income surpasses. It recommends the growth and development of individual nations because of air contamination. Fig.4 shows CEM (C-EM and C-LF), where China captured 23% of C-LF and 2% of C-EM and recorded as the highest carbon emitter, as well as Qatar, shows 28% C-EM in developing countries [15]. Germany and France trapped 9 and 6% in C-LF and, also Canada and Australia are 9 and 4% in C-EM in developed countries [5, 16–18].

**Figure 4:**
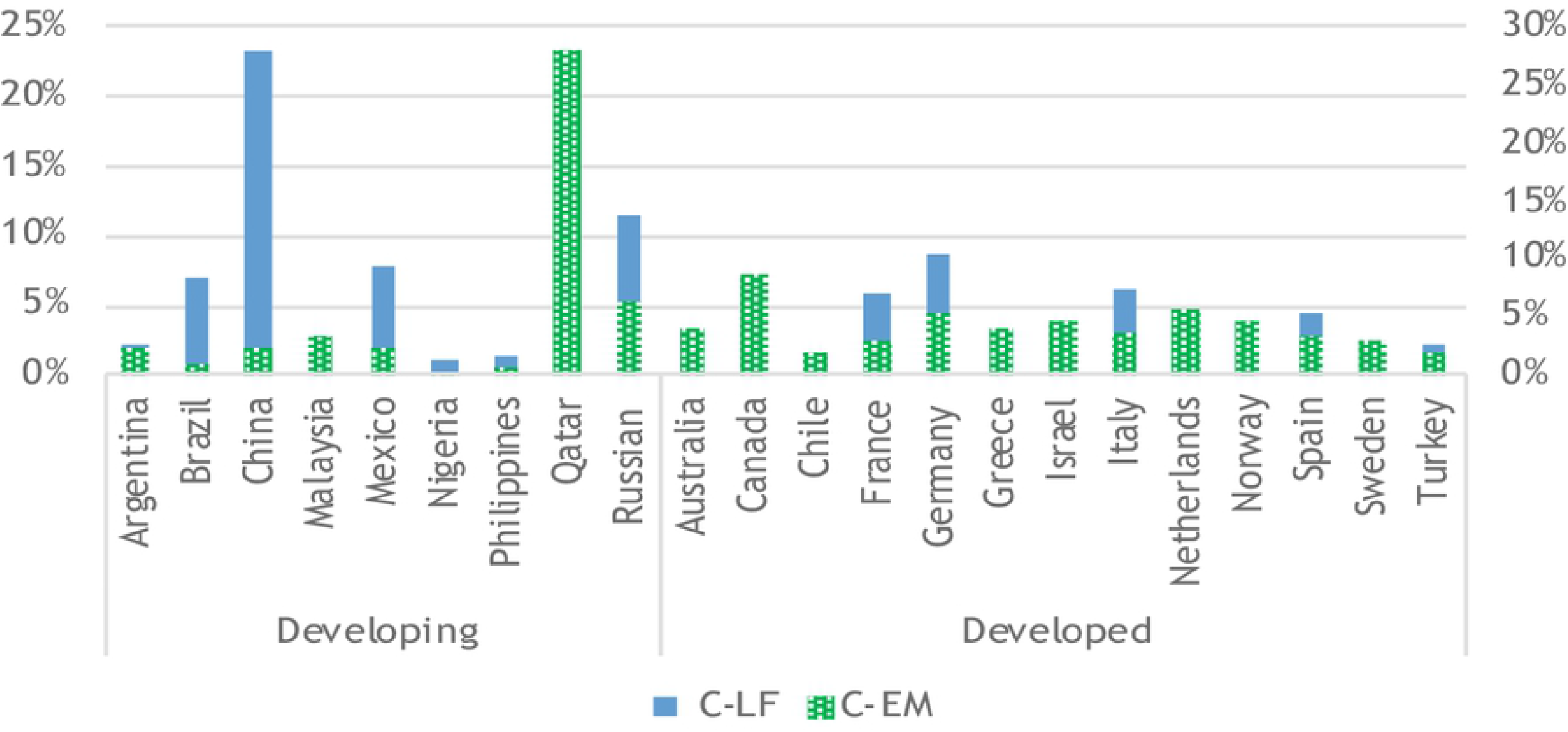
CEM of developing and developed countries.

The EKC development methodology developed presently besides of spotless afterward as well as ample serious resources besides colossal natural fetched and developing nations ought to take after the development way than that of EKC [19–21]. Within the rising economics, a significant division of the generation fulfills the utilization in developed nations because of notorious CO2 spillage issues besides encapsulated at CO2 emissions [22]. Subsequently, the developing economy like China, and Indonesia, the predicament of economic development and the same tending to environmental issues, ought to consider vitality sparing and exchanging to non-remnant fuel and carbon tax.

Based on the above mentioned, this paper offers new insights into the prior study by analyzing a variant of the EKC hypothesis-which nominates an inverted U-shape relationship between national income (while controlling for agriculture production effects) and environmental quality of 22 countries. To the best of our information, no prior study has attempted to analyze the agriculture production effects based on the EKC hypothesis in a panel of 9 developing and 13 developed countries individually. This study brings four contributions to prior studies. Firstly, we enhance the EKC model with technical effects (GNI per capita), composition effects of industry, value-added (% of GDP) by integrating agriculture (AGR), cereal yield (CY), renewable energy consumption (REN), and trade (IDC) effect on the environment. Second, due to the rapid growth of developing countries, the study utilizes two different technical models to determine the EKC effect. While the first model (total CO2 emissions-C-EM) as per endogenous variable and second model share of (CO2 emission-C-LF) by sources of liquid. Agriculture products induced CO2 emissions by agriculture mechanization and cultivation strategies in developing and developed countries. Thirdly, the panel quantile regression approach for distribution heterogeneity which includes non-additive fixed effects (FE), and it determines the CO2 emissions effects across countries [23–25]. And analyzing the exogenous indicator effects across the conditional distribution of CO2 emissions. Fourthly, the research employs panel estimation (cross-sectional dependence) techniques and finding developing and developed countries’ policy framework towards ways it can instantaneously confirm agriculture productivity while maintaining and modified clad environment in 22 countries. The structure of this study is organized in the following; Section 2 gives the review of extant literature, Section 3 shows the data collection methodology procedure, Section 4 result outlines while Section 5 concludes and recommendation.

## 2. Literature review

The prior studies on agricultural production and the Kuznets Curve (EKC) hypothesis have diversified results, and a lot of research is unidentified and examined the relationship between environmental degradation and agriculture production. And which makes no clear divergence among CO2 emissions emitting by agriculture production and emissions stimulated from the different methods of cultivation practice and new mechanical strategies. Agriculture stipulates a livelihood for millions of people in developing and developed countries every day and feeds all of us with different kinds of agriculture production. Hence, the agriculture-related product has a significant impact on environmental degradation and also a contribution to green gas emissions. According to <u>Climate Watch</u> the CO2 emissions originate from agricultural production (livestock), and more than 70 billions animal are annually raising for human consumption. The fermentation process is one of the rich sources of methane that occurs in ruminant animals (cow, sheep, cattle, and goats), and 40% CO2 emissions recorded in the last 20 years [26, 27]. Additionally, enteric fermentation, manure left on pasture, rice cultivation, synthetic fertilizers, manure management, and others (crop residues; cultivation of organic soils; burning-savanna and manure applied to soils) with 39.3%,15.2%, 10.1%,11.8%, 6.8%, and 16.8% respectively are the top causes of agriculture emissions.

Besides, Appiah analyzed the effects of crop and livestock production on CO2 emissions tended to positive but of different magnitudes. Their results indicate that a 1% increase in each agriculture production indicator increases CO2 emissions simultaneously in developed countries [28]. In 2019, Qiao examined the greenhouse effects of developing countries, that study found agricultural production activities significantly impact CO2 emissions among the developing economies, and renewable energy drops carbon emission in developed countries. There is a U-shaped of EKC in developed economies but, yet to reach its peak level in the developing countries [29]. However, this study differs essentially from others (Table 1) since it does not just analyze twenty-two countries but also distinctly segregates CO2 emission dynamics from the agriculture cultivation process and mechanization. This implication gives the government and individual entities in the developing and developed countries proper direction and policies to control emissions.

**Table 1:** Prior studies of Agriculture productivity and CO2 emissions

| Study | Datasets | Econometric techniques | Period | Outcomes |
| --- | --- | --- | --- | --- |
| [30] | G7-countries | Cross sectionally augmented auto-regressive distributive lag (CS-ARDL) | 1991-2017 | CO2 emissions reduce by green growth and renewable energy use are found to decrease emissions. |
| [9] | Developing country-China | Input and output method | 2007-2017 | China agriculture input structure effect on the growth of energy related CO2 emissions. Industrial system need to modified and reduce on China's agriculture and also supply-side. |
| [31] | 89 countries | Decoupling technique | 2000-2016 | Only 18 countries have strong decoupling and this study is analyzed the relationship between production and agriculture energy consumption |
| [32] | Malaysia | Cointegration test | 1978-2016 | Outflow of carbon emanations through Kuznets curve |
| [33] | 12 MENA countries | Panel Smooth Transition Regression (PSTR) | 1980-2015 | Relationship between GDP, CO2 emissions, and renewable electricity consumption and effect on environment by disparity in the level of income. |
| [34] | 15 ECOWAS countries | Panel quantile regression approach | 1990-2015 | Total CO2 emissions increase while agriculture production impedes by liquid sources (CO2 emissions) |
| [35] | 20 countries in Middle East and North Africa (MENA) | Regression | 1980-2014 | The existing hypothesis effect on affluence, huge technology and populace |
| [36] | India, Indonesia, China, and Brazil | GMM and Chow test | 1971-2014 | Developed countries economic complexity was creating negative impact on CO2 emissions. |
| [18] | France, Portugal and Spain | Autoregressive Distributed Lag (ARDL) | 1970- 2013 | CO2 emissions produced by agriculture and proved that different socio economic condition mitigate the greenhouse gas emissions. |
| [37] | 63 developing and developed countries | Cointegration regression | 1990-2010 | Use of renewable energy and it create positive effect on environment and economic growth. |
| [38] | Middle East, North Africa, Sub-Saharan Africa | DOLS and VEC | 1980-2010 | Existing relationship of renewable vitalities and their consumption. |
| [39] | Agriculture structure of Iran | Theil index and Kaya factor approach use to analysis. | 1960-1997 | Driving factors of agriculture industries is noticeable CO2 emissions due to inefficient policies rising fossil energy consumption. |
| [40] | 100 year timescale | Experimental (Phragmites australis and Typha latifolia) | 20 years | Global warming potential as it reduce GHG emissions from the soil. |

## 3. Data and methodology

### 3.1. Data specification

According to the <u>human development index (HDI)</u>, 25 developing and developed countries have been distributed by social and economic development levels. It quantifies attainment of education, life expectancy, and standardized income among zero and one; the most closer to one, the more developed countries, and more developed countries have HDIs of 0.8 or higher. Based on the availability of data, twenty-two countries have approached by the (<u>World Bank</u>) in this study. Under the different level of income, nine developing (Argentina, Brazil, China, Mexico, Malaysia, Nigeria, Philippines, Qatar, Russian Federation) and thirteen developed (Australia, Canada, Chile, Germany, France, Greece, Israel, Italy, Netherlands, Norway, Spain, Sweden, and Turkey) countries have analyzed and classified. The extracted database contained from 1991 through 2016 also the length of data is determined by data availability. Consistent with <u>JRC, climate change</u> 2020’s reports, and World Bank data, this study adopts two main regressor’s, notably: C-EM and C-LF are the CEM (metric tonnes per capita) although C-LF considered due to liquid consumption in individual developing and developed economies. Based on similar studies, regressors of CEM have chosen [5, 16, 36, 41–43] and individually analyzed. Due to data limitations, 25 years period consider in 22 developing and developed countries. The determinants of CEM retained explanatory indicators are coherent with prior findings [44–48].

The retained explanatory indicators include AGR, CY, REN, GNI, IDC, and TRD, which are inspecting in panel A and B Table 2. Descriptive statistics computed for each residual in the group, where the Jarque-Bera statistics reject the hypothesis of normal distribution in AGR, CY, REN, GNI, IDC, and TRD with experimental indicators provided in Table 3. In the Fig.5 industries value-added (IDC) and trade (TRD) is computed by the standard deviation in standard bars, whereas developing (Brazil, Mexico, and Russian) countries high trade value recorded, and a developed country (Spain, and Canada) specified low value-added of industries.

**Table 2:** Indicator definitions

| Indicators | Code | Sources |
| --- | --- | --- |
| CO2 emissions (metric tons per capita) | C-EM | EN.ATM.CO2E.PC |
| CO2 emissions from liquid fuel consumption (kt.) | C-LF | EN.ATM.CO2E.LF.KT |
| Agriculture, forestry, and fishing, value added (constant 2010 US\$) | AGR | NV.AGR.TOTL.KD |
| Cereal yield (kg per hectare) | CY | AG.YLD.CREL.KG |
| Renewable energy consumption (% of total final energy consumption) | REN | EG.FEC.RNEW.ZS |
| GNI per capita (constant 2010 US\$) | GNI | NY.GNP.PCAP.KD |
| Industry (including construction), value added (% of GDP) | IDC | NV.IND.TOTL.ZS |
| Trade (% of GDP) | TRD | NE.TRD.GNFS.ZS |

**Table 3:** Descriptive statistics summary

|  | **C-EM** | **C-LF** | **AGR** | **CY** | **REN** | **GNI** | **IDC** | **TRD** |
| --- | --- | --- | --- | --- | --- | --- | --- | --- |
| Mean | 8.653 | 157973.200 | 4.92E+10 | 3962.622 | 20.512 | 26006.690 | 28.976 | 63.887 |
| Median | 6.421 | 76706.310 | 2.60E+10 | 3606.800 | 11.045 | 23379.790 | 27.071 | 56.421 |
| Maximum | 70.042 | 1336398.000 | 7.16E+11 | 9073.700 | 88.832 | 93958.630 | 73.469 | 220.407 |
| Minimum | 0.312 | 2497.227 | 63721950 | 981.300 | 0.000 | 1209.466 | 13.682 | 13.753 |
| Std. Dev. | 10.607 | 193499.000 | 9.96E+10 | 1763.868 | 21.131 | 21314.980 | 8.396 | 34.420 |
| Skewness | 3.778 | 3.149 | 4.486461 | 0.660 | 1.608 | 0.902 | 1.526 | 2.015 |
| Kurtosis | 18.226 | 16.015 | 23.90555 | 2.785 | 5.095 | 3.419 | 7.409 | 8.147 |
| Jarque-Bera | 6837.624*** | 4947.522*** | 11537.17*** | 42.518*** | 337.697*** | 79.433*** | 639.753*** | 1013.352*** |
| Observations | 568 | 568 | 535 | 571 | 550 | 556 | 534 | 569 |

**Note:** Table 2 shows the definition of the indicators **Sources:** Author's estimates

**Figure 5:**
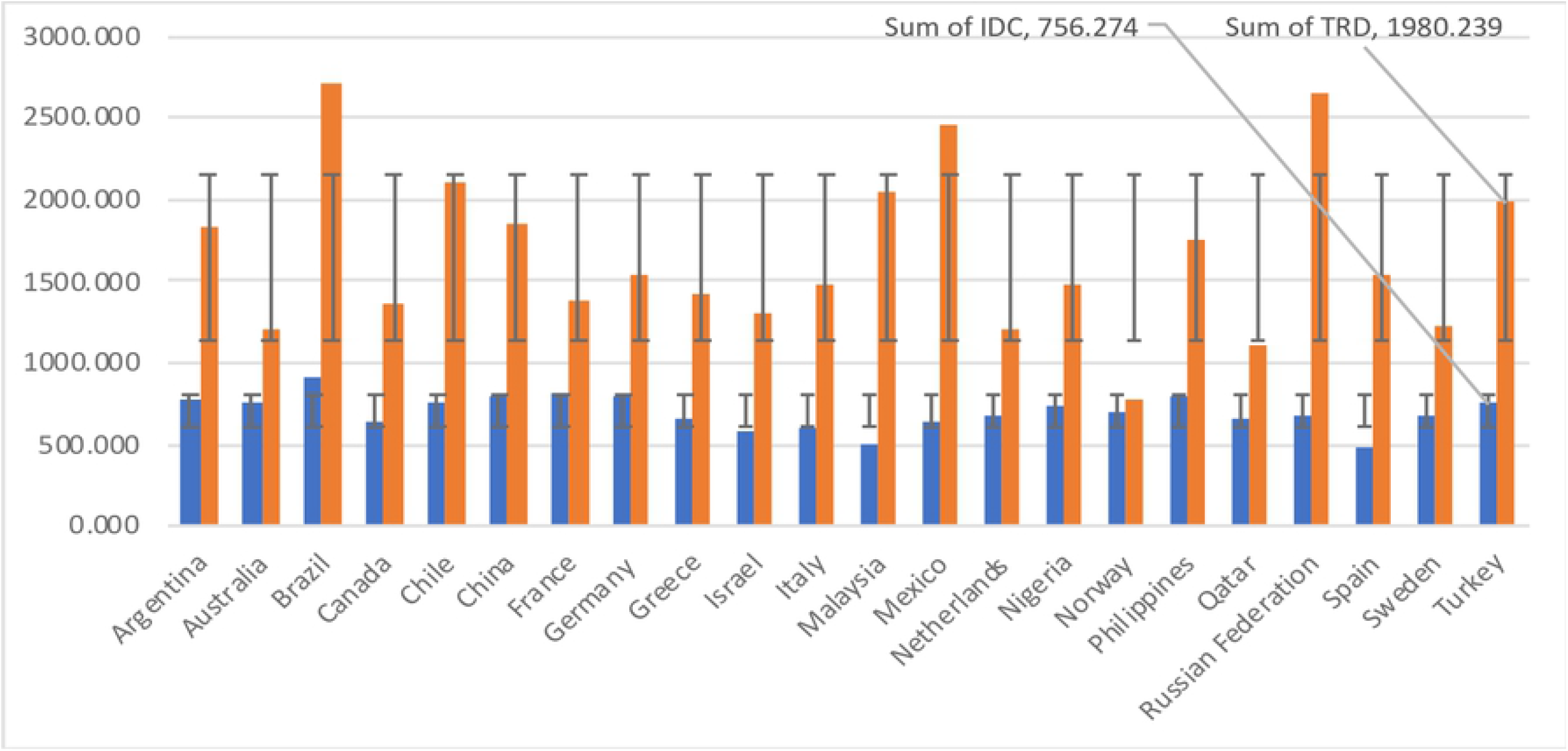
IDC and TRD pivot chat.

The panel covariance is used to examine the nature of the relationship between 22 (9 developing and 13 developed) countries. The stimulating covariance is used for analysis by an ordinary method and also computing cross-sectional (Zellner SUR-type) estimators for panel A and B in tests of cross-section dependence Table 4. In panel A, positive covariance sustained (C-EM, CY, GNI, IDC, and TRD) and panel B (C-LF, AGR, CY, and IDC), and the estimated results indicated that one indicator increases affect the other. Additionally, a negative estimation of the indicators represents an inverse relationship and always move in a different direction in panel A and B. The correlation matrix of corroborates is showing relatively diverse relationships across income levels. For instance, in the LIG countries, a positive linear relationship appears between CO2EM and GDPC, which is nonlinear in LmIG countries. Also, AGRI and CO2 emissions appear to be inverse.

**Table 4:** Covariance analysis

| Correlation | C-EM | C-LF | AGR | CY | REN | GNI | IDC | TRD |
| --- | --- | --- | --- | --- | --- | --- | --- | --- |
| C-EM | 1.000 |  |  |  |  |  |  |  |
| C-LF | 0.041 | 1.000 |  |  |  |  |  |  |
| AGR | -0.124 | 0.863 | 1.000 |  |  |  |  |  |
| CY | 0.293 | 0.161 | 0.092 | 1.000 |  |  |  |  |
| REN | -0.440 | -0.222 | 0.014 | -0.311 | 1.000 |  |  |  |
| GNI | 0.534 | -0.220 | -0.292 | 0.399 | -0.020 | 1.000 |  |  |
| IDC | 0.243 | 0.283 | 0.363 | -0.122 | 0.048 | -0.206 | 1.000 |  |
| TRD | 0.251 | -0.210 | -0.170 | 0.264 | -0.221 | 0.135 | 0.322 | 1.000 |

**Note:** Table 2 shows the definition of the indicators. **Sources:** Author's estimates

### 3.2. Methodology

The description of the model is showing as:

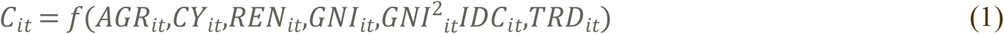

Where *C_it_* denotes carbon dioxide emission for the two dependent indicators (C-EM and C-LF). AGR indicates the agricultural value-added includes forestry, fishing, hunting, livestock production, and cultivation of crops. CY shows cereal yield, the measure by harvested land, includes wheat, maize, rice, etc. REN represents the consumption of renewable energy. *GNI^2^* denotes gross national income, which is divided by the midyear population and captures the nonlinear effect of per-capita income on environmental quality. IDC shows industry (value-added) corresponds to the International Standard Industrial Classification (ISIC). TRD implies trade (export and import) goods and services measure as a share of GDP. And *t* and *i* the time and country dimensions of panel data in Table 1. In the spirit of the Kuznets curve (EKC) model, Eq. (1) is showing as follows:

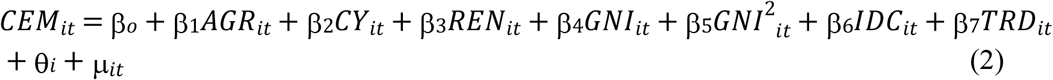

Eq. (2) indicated the coefficients of ßs that capture the influence of covariates on responding indicators (*CEM_it_*) while *θ_i_* and *μ_it_* are the state fixed effects and the stochastic error term having a zero mean with constant variance. The panel data estimated by Random Effects (RE) and Fixed Effects (FE) as well as Fully Modified OLS (FMOLS) and Dynamic OLS (DOLS) estimators [49, 50] that create asymptotically unbiased, normally distributed coefficient estimates. The presence of cross-section implies fixed and random effects, and restrictions stipulate a model containing impact in both (cross-session and period) dimension. EKC and inverted U-Shaped relationship between environmental quality and output. The estimated value of *β*_4_ and *β*_5_ and are respectively likely to be positive and negative through the estimators.

### 3.3. Panel quantile regression

Based on the priors studies, conditional determinants of CO2 emissions are the essential majority of this model and are concerned with examining the conditional mean of analysis (C-EM and C-LF) indicators across countries. The increasing approach of this model to quantiles and provides a linear relationship between repressors in panel A and panel B. As for unbiased estimations [51], we used quantile regression (QR) in the presence of potential outliers in panels. Henceforth, the QR methodology allows us to capture CEM factors with as sturdy importance on developing and developed economies in environmental degraders, which are high in developing countries comparatively developed [27, 51].

Relatively, more income and secure economics tend to be extra diligent, and from the growth, an extension has more influence on the quality of the environment. Hence, there are potential heterogeneities across developing and developed economies by environmental degradation. Moreover, the developing economies are highly involved in agrarian, and the territory of the environment depending on agriculture practice.

Unlike the conventional QR, this study adopts by [52] maintains a panel data framework by non-additive fixed effects with the term of inseparable disturbance. Accordingly, this research specification allows the repressors to be non-separable from developing and developed countries’ fixed-effects. Therefore, the panel quantile specification is as follows.

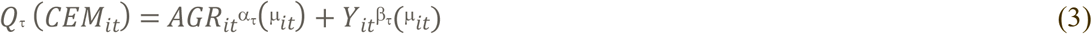

Eq.(3) indicated that *Q_τ_* as the conditional quantile of CEM such that τ = (0.25,0.50, and 0.75). hence, individual country heterogeneous effects *μ_it_* = *f*(Ø_*i*_,∂_*it*_) are inextricable from indicators in the panel and AGR is as stated while Y capture further co-variates AGR, CY, REN, GNI, IDC and TR

### 3.4. Quantile decomposition

As for decomposition of developing and developed countries data in the predictive performance of quantile and for well-detailed analysis of individual state to determine the impact of agriculture and their influence on environmental degradation by an economic factor on CEM (C-EM and C-LF), where this study will further examine the extent of panel A and B in developing and developed economies by Low Income Group (LIN_G) and High Income Group (HI_G). The developing and developed countries were containing $6,376 to $37,760, with a total population of 5.74 to 1.30 billion in 2012. That segment allows us to investigate the extent to which CEM different levels of income in panels A and B. This decomposition approach in our research study is showing economies due to [25, 53–55]. In this current context, we use the quantile decomposition test in panel A and B [56–58]. The CEM decomposed by decomposition techniques of LIG and HIG into two modules like as income groups of developing countries differences in perceived agriculture or income factors, and income group’s difference in return across economies in developed countries and minimum environmental degraders.

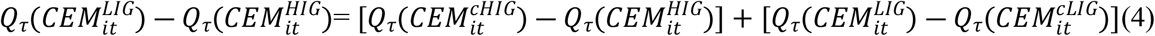

Eq.(4) is indicating 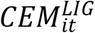 and 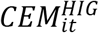 the regressors of CEM for LIG, and HIG also shows the distribution of counterfactual. Which compares the quality of the environment should possess economic characteristics and agriculture as the LIG in developing economies. The quantile regression captures by *Q_τ_* (Q1,Q2, and Q3). Like prior research, there are different levels of income capture in agriculture and economic factors in the developing and developed countries.

## 4. Finding and results

### 4.1. Panel unit root test

The unit root test is evaluated the hypothesis of stationary (trend) by analyzing whether the absolute value of ρ is sternly less than one in panel A and B. It generally specified the null hypothesis (*H_0_*:ρ = 1) beside the one-sided alternative (*H_0_*:ρ < 1). The unit-root test is investigated and explore the intercept level and trend in panels A and B. Prior studies to estimating the regression parameters and properties of the indicators examined to delineate which empirical method would best suit the analysis. We use the panel root test and, it allows for heterogeneous autoregressive coefficients ADF [59–61].

The persistence parameters analyzed by common across (CA) or freely across (FA) cross-sections, where all the tests LLC, Breitung, and Hadri employ by CA and IPS, Fisher-ADF, Fisher-PP used FA Table 5. A chi-squared statistics of the ADF principle studied in such a way that the ρ-values individually specify countries (Augmented) test and summed across panel members by a χ-squared distribution with 2 N d.f under the null of no stationary in the panel series.

**Table 5:** Unit root test (Level and 1st difference)

| Individual intercept<br>(Level) |  |  |  |  |  | Individual intercept and trend<br>(1<sup>st</sup> difference) |  |  |  |  |  |
| --- | --- | --- | --- | --- | --- | --- | --- | --- | --- | --- | --- |
| Var | CR | Individual root |  |  | Hadri | CR |  | Individual root |  |  | Hadri |
|  | LLC | IPS | ADF | PP |  | LLC | Breitung | IPS | ADF | PP |  |
| **C-EM** | 1.702** | 2.910** | 21.044** | 56.938** | 9.377*** | -2.450*** | -3.958*** | -6.927*** | 128.413*** | 570.882*** | 8.245*** |
| **C-LF** | 1.603** | 1.597** | 42.221** | 46.093*** | 16.868*** | -4.188*** | -2.357*** | -8.574*** | 157.564*** | 621.065*** | 10.510*** |
| **AGR** | 3.102** | 5.010* | 24.684** | 30.987** | 18.206*** | -5.4075*** | -3.388*** | -6.671*** | 150.017*** | 1048.54*** | 7.302*** |
| **CY** | -0.0581** | 1.552** | 36.141** | 79.567*** | 15.581*** | -6.1331*** | -3.098*** | -13.737*** | 241.143*** | 2874.28*** | 6.746*** |
| **REN** | 1.277** | 2.567** | 37.246** | 48.175*** | 14.364*** | -8.833* | -5.568*** | -9.011*** | 157.316*** | 673.611*** | 7.549*** |
| **GNI** | -0.458** | 3.622** | 24.549*** | 26.678** | 16.666*** | -7.798*** | -3.762*** | -6.570*** | 128.692*** | 204.653*** | 7.993*** |
| **IDC** | -2.369*** | -0.360** | 53.112** | 62.153*** | 10.189*** | -10.775*** | -5.950*** | -9.412*** | 164.903*** | 312.410*** | 9.765*** |
| **TRD** | -3.969*** | -2.766*** | 76.019*** | 73.893*** | 9.916*** | -8.427*** | -9.931*** | -9.974*** | 172.577*** | 413.410*** | 9.203*** |

**Note:** Table 2 shows the definition of the indicators. **Sources:** Author's estimates Individual effects of exogenous variables have been computed by user-specified lags 1 in the Newly-West automatic bandwidth selection and Bartlett kernel. The probabilities for Fisher tests as asymptotic square distribution. \*\*\* denotes significance at the 1% level, \*\* at the 5% level, and \* at the 10% level. Levin, Lin, and Chu (LLC), The Im, Pesaran, and Shin (IPS) Augmented Dickey-Fuller (ADF) and Phillips-Perron (PP) Tests

The existence of a single latent factor controls by the IPS test allows for heterogeneity in the autoregressive coefficient (ac) of the ADF regression, and this mitigates incidences of cross-sectional dependence in the data. It also contains an average cross-section of regressors and explanatory indicators. A non-standard distribution follows under the null hypothesis of non-stationary in panels A and B. The unit-root test shows that individual intercept by level and 1^st^ difference and implying that indicators integrated of order 1, or I(1) even after controlling for both intercept and trend terms.

### 4.2. Panel cointegration test

After proving the non-stationary of all the indicators in both models. It becomes necessary to determine if the variables are cointegrated [49, 50]. Padroni panel-test by a statistical technique (Kao) investigated, where the panel B cointegration employed with ADF, PP, and rho statistics, and the test results indicated that 2.489 with dynamic residual value Table 6. The test result highlighted those non-stationary indicators have a high potential of being spurious in panel A and B models. Additionally, the statistical test of Fisher, that dynamic methodology expanded to the causal method of Johansen by panel cointegration test [62], presented in Table 7. The estimation results of cointegration gather with the identical value of the p-value of the separated statistics of the Johansen test [63]. And it rejects the cointegration of the null hypothesis.

**Table 6:** Padroni (Engle-Granger based) Test

| **Panel A: Within-dimension** |  |  |  |  |  |  |  |  |
| --- | --- | --- | --- | --- | --- | --- | --- | --- |
| Panel co-integration test | Carbon emissions (C-EM) Equation |  |  |  | Carbon emission from liquid (C_LM) Equation |  |  |  |
| Common AR | Individual intercept |  | Individual intercept and trend |  | Individual intercept |  | Individual intercept and trend |  |
|  | Statistic | Weighted Statistic | Statistic | Weighted Statistic | Statistic | Weighted Statistic | Statistic | Weighted Statistic |
| Panel v-Statistic | -0.305** | -0.886** | -2.094** | -2.358** | -1.498** | -2.867** | -3.316** | -4.469* |
| Panel rho-Statistic | 2.768** | 2.364** | 4.368* | 3.760** | 2.821** | 1.977** | 5.376* | 3.769** |
| Panel PP-Statistic | -0.518** | -2.630** | 1.087** | -2.257** | -2.809*** | -7.838*** | 0.172** | -9.018*** |
| Panel ADF-Statistic | -0.180** | -0.200** | 1.024** | 0.348** | 1.057** | -2.799*** | 4.258** | -2.471*** |
| **Panel B: Between- dimension** |  |  |  |  |  |  |  |  |
| Individual AR | Statistic of individual AR coefs. |  |  |  |  |  |  |  |
|  | Individual intercept |  | Individual intercept and trend |  | Individual intercept |  | Individual intercept and trend |  |
| Group rho-Statistic | 4.067* |  | 5.004** |  | 4.162* |  | 5.844* |  |
| Group PP-Statistic | -2.976*** |  | -3.249*** |  | -7.369*** |  | -11.688*** |  |
| Group ADF-Statistic | 0.868** |  | 1.699** |  | -1.020** |  | -1.130** |  |

**Note:** Table 2 shows the definition of the indicators. **Sources:** Author's estimates. Criterion of Info Schwarz selected the lag length. \*\*\* stipulate at the level of 1%\*\* stipulate at the level of 5% \* stipulate at the level of 10%

**Table 7:** Johansen Fisher Panel Cointegration Test

| Hypothesized No. of<br>CE(s) | **Carbon emissions (C-EM)**<br>**Equation** |  | **Carbon emission from liquid**<br>**(C_LM)Equation** |  |
| --- | --- | --- | --- | --- |
|  | Fisher Stat.*<br>(from trace test) | Fisher Stat.*<br>(from max-eigen<br>test) | Fisher Stat.*<br>(from trace test) | Fisher Stat.*<br>(from max-eigen<br>test) |
| None | 790.5*** | 700.2*** | 807.5*** | 789.9*** |
| At most 1 | 528.9*** | 328.9*** | 581.5*** | 339.1*** |
| At most 2 | 349.9*** | 230.3*** | 397.2*** | 281.9*** |
| At most 3 | 276.7*** | 173.1*** | 271.5*** | 163.0*** |
| At most 4 | 146.9*** | 82.66*** | 145.1*** | 87.21*** |
| At most 5 | 104.0*** | 76.12*** | 95.51*** | 84.91*** |
| At most 6 | 90.35*** | 90.35*** | 61.32*** | 61.32*** |

**Note:** Table 2 shows the definition of the indicators. **Sources:** Author's estimates. Probabilities selected by lag (11) and Criterion of Info Schwarz standard computed using asymptotic Chi-square distribution. \*\*\* stipulate at the level of 1%\*\* stipulate at the level of 5% \* stipulate at the level of 10%.

### 4.3. Panel regression

Table 8 is indicating the outcome of fixed and random influence as well as the FMOLS regression for each dependent indicator (C-EM and C-LF) in panels A and B. The FE and RE regressions effect were examined and analyzed with standard error, which is robust to occurrences of cross-sectional influence [64, 65]. The FE estimator controls for individual intercepts by Fully Modified OLS, and it has added advantage to observing for heterogenous serial correlation properties in the panel data via a non-parametric process. The FE and RE are discriminated by the Hausman test, which shows that the FE is well-effectual to the other specific model. Accordingly, the analysis will rest on the FMOLS by FE estimation, and RE reported for comparative purposes.

**Table 8:** FMOLS, Random Effect (RE), Fixed Effect (FE), Quantile regression

| Variables | FMOLS |  |  |  | Fixed effect (FE) |  | Random effect (FE) |  | Quantile regression |  |  |
| --- | --- | --- | --- | --- | --- | --- | --- | --- | --- | --- | --- |
|  | Coefficient covariance matrix |  | Residual diagnostic | Coefficient diagnostic |  |  |  |  | 1<sup>st</sup> Quarter | 2<sup>nd</sup> Quarter | 3<sup>rd</sup> Quarter |
|  | Default (Hom-V) | Sandwich (Het-V) | AC | Wald Test | Ordinary | White CS | Ordinary | White CS | Q25 | Q50 | Q75 |
| **Panel A: Carbon emissions (C-EM)** |  |  |  |  |  |  |  |  |  |  |  |
| AGR | 6.188*** | 22.464*** | 0.637 | 6.189*** | -19.129*** | -2.391** | -5.923*** | -1.130** | 1.160** | 2.027*** | 2.925*** |
| CY | -1.974** | -3.144*** | 0.392 | -1.974** | 1.383** | 0.238** | 0.263** | 0.054** | -0.481** | 1.131** | -2.823*** |
| REN | -10.625*** | -22.603*** | 0.194 | -10.625*** | -76.284*** | -12.352*** | -29.537*** | -5.205*** | -7.731*** | -10.766*** | -9.996*** |
| GNI | 5.551*** | 7.360*** | 0.138 | 5.551*** | 87.793*** | 13.423*** | 34.554*** | 7.709*** | 20.051*** | 23.592*** | 7.798*** |
| IDC | 3.109*** | 5.355*** | 0.114 | 3.109*** | 66.000*** | 3.991*** | 24.869*** | 2.791*** | 1.139** | 0.107** | 0.125** |
| TRD | -2.932*** | -4.250*** | 0.045 | -2.931*** | -15.782*** | -2.071*** | -6.321*** | -1.249*** | 1.254** | 3.623*** | 1.439** |
| Constant | - | - | - | - | -10.004*** | -0.759** | -4.402*** | -0.980** | 3.287*** | 5.834*** | 8.844*** |
| **Panel B: Carbon emissions from liquid (C-LF)** |  |  |  |  |  |  |  |  |  |  |  |
| AGR | 31.140*** | 42.456*** | 0.673 | 31.139*** | 165.080*** | 36.757*** | 33.680*** | 7.814*** | 19.417*** | 41.602*** | 14.143*** |
| CY | -0.465** | -0.890** | 0.530 | -0.464** | 8.267*** | 1.647** | 1.294** | 0.264* | 2.560*** | 1.567** | -0.249** |
| REN | -3.877*** | -6.572*** | 0.371 | -3.876*** | -59.901*** | -13.944*** | -12.313*** | -2.538*** | -7.932*** | -6.581*** | -9.554*** |
| GNI | -0.250** | -0.590** | 0.229 | -0.250** | 7.014*** | 2.106*** | 1.536** | 0.307** | 4.460*** | 2.660*** | -0.004** |
| IDC | 6.296*** | 8.720*** | 0.136 | 6.296*** | 12.315*** | 1.696*** | 2.997*** | 0.547** | 2.254*** | 0.694** | 1.080** |
| TRD | -0.183** | -0.337** | 0.032 | -0.182** | -33.038*** | -7.651*** | -7.339*** | -1.521** | -3.073*** | -2.983*** | -6.234*** |
| Constant | - | - | - | - | 31.588*** | 4.936*** | 4.960*** | 1.055** | 0.065** | 2.669*** | 2.695*** |

**Note:** Table 2 shows the definition of the indicators. Homogenous variance (Hom-V) and Heterogenous variance (Het-V). **Sources:** Author's estimates. Panel A-weighted statistics value of R-square has been computed individually for FE and RE with 0.629 and 0.706. The Durbin-Watson stat is 0.057 and 0.014. Panel B, R-square of FE and RE respectively 0.817 and 0.913 and the Durbin-Watson stat are 0.030 and 0.004 with observation (n=11385) in 23 cross-sections. The coefficient covariance method of random effect has computed by White cross-section and weighing by Swamy-Arora. The residual diagnostic of indicators is obtained from the correlogram test and AC (6 lag specification). \*\*\* denotes significance at the 1% level, \*\* at the 5% level, and \* at the 10% level.

The estimated coefficient of the panel (A and B) are accessible in the middle of the output. Of central importance is the coefficient’s on AGR, CY, REN, GNI, IDC, and TRD, which denotes that the expected Cointegrating Vector (CV) for C-EM and AGR, CY, REN, GNI, IDC, and TRD. The standard error, t-statistic (p-value) have been computed individually for C-EM and C-LF. First, all of the fit statistics are analyzed using the raw data, not the transformed data of FMOLS. The Nonstationary has used pooled estimation, and it performs standard FMOLS on the pooled sample after eliminating the deterministic components from both the experimental indicator and the regressor. As for Pooled FMOLS estimation, the long-run covariance is computed by Default (homogenous variance) and a Sandwich (heterogeneous variance) in coefficient covariance matrix with d.f. Adjustment, presented in Table 8. All indicators in panel A and B statistically significant and are robust to estimators employed.

The C-EM and C-LF estimated results for both equations in panels A and B uncovers some latent pragmatic dynamics. The indicated results of the EKC hypothesis is valid in the 22 (9 developing and 13 developed) countries [27, 66]. However, the C-LF presence is higher in the developing countries and employed to all robust results. The GNI is showing a positive relationship with 5.551 in the coefficient of the covariance matrix in panel A. However, GNI results indicated a negative relation with −0.250 in panel B.

The outcome of AGR has a positive and significant influence on C-LF, implying that a 1% change in AGR is associated with a 36.75% increase in environmental degradation. In the developing countries, agriculture-related activity influence negatively environmental problems, likewise deforestation for feed cropping, burning of biomass, and deep soil cropping [16, 29, 58, 67]. Alternative influence of the C-EM measure of environmental degradation shows that the AGR indicator is negatively related, and this means that a 1% change in AGR decreases the share of carbon emission in total emissions by 19.12%. The results show that it does not imply a reduction in the overall carbon emissions in developing and developed countries but a reduction in the mechanization of agriculture induced emissions.

It may also imply the industrial (IDC) sector by fuel-intensive of developing and developed economies due to agriculture exports [36, 37, 68]. The inverted U-shaped association detected across developing and developed economics [16, 69]. Although, this might be insinuating that the developing economies are on the economic growth level rather than ecological quality in the short-run and sustainable environment technique affect in the long-run. Additionally, environmental degradation mitigates by the adoption of renewable energy (REN) using both indicators CEM, the industrial revolution. However, the environment degrades precautions through an increase in the C-LF caused by the high consumption in developing countries. And, TRD openness decreased in both panels. For instance, a 1% change in trade openness decreased in C-EM and C-LF with 15.78% and 33.03%. These results indicated that economic structure and nature of trade of developing and developed economies [37]. Trade may also improve the environmental quality precaution through techniques and transfer clean technology. This increase in trade openness in especially developing economies with environmentally friendly policies with ideal environmental implications [70].

While measures of fit and the Durbin-Watson of FE (0.057) and RE (0.014). The CV is (1,-1) and Coefficient Restrictions (CR) with “C(1)=0 is estimated output results of Coefficient diagnostic (CD)/ Wald test. The estimated result of t-statistics (p-value) is around 0.000, indicating that we rejected the null hypothesis of the panel (A and B) that the coefficient value of the cointegrating regressor is equal to 1 with linear restriction. The Correlogram-Q-statistics reported by the residual diagnostic value of Akaike AC’s (−2(l/T)+2k/T) is estimated (at lag 6, d.f.=6) where the d.f.= the number of lags. For lags 1 to 6, the AC’s are within the ‘error bars’ infers that the p-values of panel A, where the TRD is less than 0.05 (at 0.045) and in panel B (at 0.032) therefore, residual are white noise and we can reject the null hypothesis. We estimated the pool equation by the Fixed Effects (FE) and Random Effects (RE). In the FE and RE, a model regressing (panel A and B) is analyzed by regressors (AGR, CY, REN, GNI, IDC, and TRD) and using cross-sectional identifiers only for the FE.

The coefficient covariance is analyzed further by default and White cross-section”. The Coefficient Standard Errors (CSE) and Robust Coefficient Covariance (RCC) are judges from the default and White cross-section. The Effects Specification (ES) is measure by cross-section, period, and idiosyncratic error components of S.D and Rho in FE and RE. The intraclass correlation or Rho gives that proportion of variation in the regressor (0.000).

### 4.4. Quantile regression

The quantile regression (QR) estimator is the least absolute deviations (LAD), which relates to appropriate the median of response indicator. It is a more ample analysis technique of the conditional distribution than conditional mean analysis alone. Panel A and B are indicating the 1^st^, 2^nd^ and 3^rd^ quartiles. Where the 25^th^, 50^th^, and 75^th^ percentile of response indicators and that are influence by regressor in the model. Additionally, the QR approach does not necessitate a strong distributional assumption. It deals with a robust method (RM) of forming these interactions. This section findings while equally addressing the limitation of explanatory indicators results of panels A and B. We also report the heterogenous determinants of environmental degraders in Table 8 by 25^th^, 50^th^, and 75^th^ percentiles. The conditional determinants estimates of panel A and B equations vary along the quantiles representing the heterogeneity of determinants of environmental degradation across the developing and developed countries. In panel A and B total equations, the impact of AGR on C-EM and C-LF is heterogeneous and significant across Q25, Q50, and Q75 quartiles. Accordingly, AGR contributes to environmental degradation, and the impact of real income on C-EM and C-LF observed to change sing with Q25, Q50, and 75 quartiles. Thus, while the EKC greatly effect is observed at Q50 and Q75, relatively Q25 less effective, but the EKC is validated.

Descriptive statistics result reported under Fig 1 above shows that Canada and Qatar are amongst the highest carbon emitters developed and developing countries wherein the EKC hypothesis validated for Nigeria, Philippines, Brazil, Chile, and Argentina. For the highest per-capita carbon emitters such as Canada, Qatar, Russian, and the Netherlands are in the upper Q50, the effect of real income of these developing and developed economics mirror the inverted U-shaped pattern of the Kuznets Curve (EKC) hypothesis.

Considering the C-LF consumption, the quantile results for all indicators despite the greatest influence and has significant pollution abatement effect except CY and IDC in Q50, where the cereal yield and industries association have little bit low effect presented in Fig 1 such as France, Italy, Norway, and Germany. Additionally, AGR, REN, and TRD reveal almost the identical influence across all quantiles as mean-based estimators.

### 4.5. Discussion

The quintile decomposition analysis for the CEM emissions shows undetected issues outside the research model tend to provoke carbon emissions gap among LIN_G and HI_G with the implication of different developing and developed economic structures and frameworks of environmental policies. However, C-LF and C-EM tend to be very high relative to the observed economic factors and agriculture. The estimated results entail that developing countries are more homogenous as regards growth strategies and changes to C-EM. Additionally, the EKC type effect for the CEM with total and liquid in panel A and B. The empirical analysis of this study point out variety of environmental degradation by agriculture production, economic growth, trade openness, Renewable energy, industry, and trade inferences in more detail below;

1. According to FAO (Food and Agriculture Organization of the United Nations) estimated that emissions from forestry, agriculture, and fisheries have nearly twice over the past 50 years and continuously increase an additional 30% by 2050. The overall energy demand rapidly increases in developing and developed countries, which means that CO2 emission is still rising and reached an all-time global high for the 4th consecutive year [71]. Consider that the Asian region only accommodates 4.3 billion people, which is 60% of the world population. The 1% increase in population in that developing countries are slowly increasing their per capita of CEM emissions have driven to more than double developed countries. In developing countries, a balance needs to maintained mechanization of CEM from unsustainable cultivation strategies, which require more addable and exactness to be more ecological [72]. GR production shows a positive effect on all Q25, Q50, and Q75 in panels A and B with an EKC high effect. However, developing and developed countries still need modern methods of agriculture production practiced by the low and high levels of income in rural and urban areas. Instinctively, it can state that this implication may account for variation in CO2 emission inducing effect of agriculture.
2. The existence of the EKC hypothesis for CEM does not hold for the developing countries, and the estimated results are inconsistent with what economies obtained in the level of high income. However, empirical estimation of quantile regression indicated that the hypothesis of EKC for CEM holds at higher emission quantiles, wherein the EKC hypothesis validated for Qatar, Canada, and China relatively higher emitting economy in developing and developed countries [73–75]. The income measure (GNI per capita) links with CEM in panels A and B as observed from the FMOLS model, and identified the EKC from the total and liquid carbon emissions. A variety of other factors can also play influential roles, likewise, as the pattern of heterogeneity of gaseous and solid fuel utilization in all countries. For insistence, of all the 22, countries only Qatar, Canada, and China have CO2 emissions emanating throughout the study. Others like Nigeria and Chile in developing and developed countries report fewer emissions on C-LF and C-EM Fig.4. And this may be why the Kuznets curve effect not observed for CEM, also indicated that the fact that environmental policy formation in the countries.
3. The estimated value of Trade (TRD) indicated a negative impact on panels A and B. And intensive trade has an enervating influence on the energy obtained from industrial sectors from the developing countries [76, 77]. However, developing countries highly dependent on agricultural products and primary sector exports, it is estimated that agriculture and trade would track similar dynamics in developing and developed countries. Additionally, economies balance economic growth and environmental sustainability.
4. The renewable energy throughout negative in panel A and B, looking at the quantile estimates, the obtained results at Q25 has a distinct non-standard and favorable in both panels. Renewable energy has a negative relationship with C-EM, and C-LM and a positive relationship have obtained with economic growth in the long run in developing countries [78, 79]. Thus renewable energy interacts with carbon emissions from liquid fuel consumption to negative stimulus human development and economies growth in developing and developed countries. However, the result of panel B relatively more homogenous as CO2 emissions is detected negative dynamics and robust to all mean estimators.
5. Positive and significant effect of industry value-added observed in both panels A and B. These estimations may imply that industrial sectors of developing economies may be weaker compared to developed economies. The effect of C-LF becomes higher significant (p<.1) at quantile Q25, so the energy consumption of developing countries increased by technological progress [80]. The industrial revolution is playing a vital role in economic growth. The negative effect of trade in Q25-75 in panel B indicated the influence of liquid carbon emissions in economies. The industries and trade coefficients show that agriculture and trade consequences with relatively more statistical evidence than industry [81].

## 5. Conclusion and recommendation

This study is considering the consequence of agriculture (AGR), cereal yield (CY), renewable energy consumption (REN), GNI, industries (IDC), and trade (TRD) in developing and developed countries. Environmental degradation is investigated by two sources of CEM (C-EM and C-LF) and using panel data from 22 (9 developing and 13 developed) for the period 1991-2016. This study adopted panel quantile decomposition techniques with FMOLS to explore if the relationship between AGR and economic factors with different levels of income and CO2 emitter, also the extent of the CO2 emissions gap between low (LIN_G) and high (HI_G) economies. Estimated results depict heterogeneity of environmental quality determinants in developing and developed countries, which vary through a high and low level of CO2 emitters. As an estimated outcome of the abovementioned, the study recommends the following guidelines. First, the selected developing (China, Brazil, and Nigeria) developed (France, Italy, and Spain) countries need to play a vital role in exterminating crude agriculture practices relating to the natural fragmentation of land. And giving rise to agricultural products and bush burning that have enervating environmental effects. Developing countries need more agricultural practice by cultivation techniques and apply new sustainable methods. Additionally, developed economies should utilize light and low energy equipment for the protection of the environment and also the elimination of fossil fuel-based peats. As for environmental degradation, the implementation of sustainable agriculture should protect potential household-related CO2 emissions, and in most developing countries, the layman involves the health risk issues cause by biomass waste products.

Second, government implementing strategies should be empowered and established to prevent and reduce CO2 emissions risk in urban areas in developing countries. And the dynamic modification of the green techniques to protect soil, green gas emissions, and cut energy consumption. Third, there is a need to improve precision agriculture techniques and introduce the AgriCare conservative agriculture approach, based on strip cropping and no-till farming in developing countries. Private and public sectors’ intentions bringing positive change in agriculture by natural methods and promote more sustainable economies. Fourth, modification of renewable energy innovations should be encouraged in developing countries, and environmental degradation can be controlled and protect with energy-efficient techniques and natural methods. It would enable a more sustainable transition in high polluted industrial-based economies in developing countries.

Finally, agricultural and industrial products in developing and developed countries should be balanced to cum adequate CO2 emission control and solve the main problems, which are low industrial capacity, food shortage, and untenable manufacturing practice.

